# Dissecting the Role of the Lateral Entorhinal Cortex in Memory Interference

**DOI:** 10.64898/2026.06.05.730409

**Authors:** Omar Ghazy, Manal Mansour, Jacquelyn N. Tomaio, Susana Mingote

**Affiliations:** Advanced Science Research Center at the Graduate Center, CUNY, New York, NY, USA

## Abstract

Memory interference occurs when an older memory competes with a newer memory that shares similar features ^1^. The hippocampal–entorhinal system is essential for memory ^2,3^, and the entorhinal cortex has been implicated in interference using the latent inhibition paradigm ^4^. In latent inhibition, prior non-reinforced exposure to a stimulus reduces the conditioned response later elicited when that same stimulus is paired with an aversive or appetitive outcome ^5^. Competition-based models propose that latent inhibition arises because the memory that the pre-exposed stimulus was inconsequential competes during retrieval with the memory that the same stimulus predicts an unconditioned stimulus ^6^. Although entorhinal cortex lesions or inactivation impair latent inhibition ^7–10^, the specific contribution of the lateral entorhinal cortex (LEC) remains unclear. The LEC provides non-spatial and cue-related input to the hippocampus ^11^ and has been implicated in associative memory ^12,13^. LEC function also declines with age ^14,15^, a period when memory interference increases ^16^. Here, we used chemogenetics to transiently inhibit excitatory neurons in the LEC during retrieval in a latent inhibition paradigm. LEC inhibition did not impair conditioned fear responses to a non-pre-exposed tone, indicating that retrieval of the tone–shock association remained intact. However, LEC inhibition attenuated latent inhibition by increasing conditioned fear responses to a pre-exposed tone. Together, these findings suggest that the LEC supports retrieval of prior inconsequential stimulus memories that compete with newer conditioned associations, providing a potential mechanism by which age-related LEC dysfunction may contribute to increased memory interference.

**Highlights:**

- Pre-exposure to a tone attenuates retrieval of a tone-evoked conditioned fear response, revealing latent inhibition
- Lateral entorhinal cortex inhibition does not impair fear-memory retrieval in non-pre-exposed animals
- The lateral entorhinal cortex is required for the expression of latent inhibition during retrieval
- The lateral entorhinal cortex contributes to memory interference during retrieval of stimuli with conflicting associations

**In Brief:** Ghazy et al. show that the lateral entorhinal cortex (LEC) is required for latent inhibition during retrieval, in which prior tone exposure attenuates later fear responding to that same tone. Because LEC inhibition does not impair tone-evoked conditioned fear responding, these findings indicate that the LEC contributes to memory interference when the same stimulus has conflicting associations.

## RESULTS

### Tone Pre-Exposure Attenuates Tone-Evoked Fear Responses and Produces Latent Inhibition

We first investigated whether prior exposure to a tone interferes with the subsequent expression of tone-cued fear conditioning, thereby producing latent inhibition. To test this, we used a paradigm adapted from Mingote et al. (2017) ^17^ consisting of three phases: pre-exposure, conditioning, and retrieval (**Fig. 1A**). During pre-exposure, mice in the pre-exposed (PE) group received 20 presentations of a 1400 Hz, 80 dB tone (30 s duration) per session for 3 consecutive days, delivered at random interstimulus intervals to establish the tone as behaviorally inconsequential. In contrast, mice in the non-pre-exposed (NPE) group received an equivalent number of light stimuli and were not exposed to the tone. During conditioning on day 4, the same tone co-terminated with a 2 s foot shock (0.7 mA), and this tone–shock pairing was repeated three times in both groups. During retrieval on day 5, animals were exposed to a 4-min baseline period followed by a 6-min continuous tone presentation to assess tone-evoked freezing. Critically, retrieval was conducted in a context distinct from that used for pre-exposure and conditioning to minimize contextual fear and isolate tone-evoked responding.

**Figure 1.**
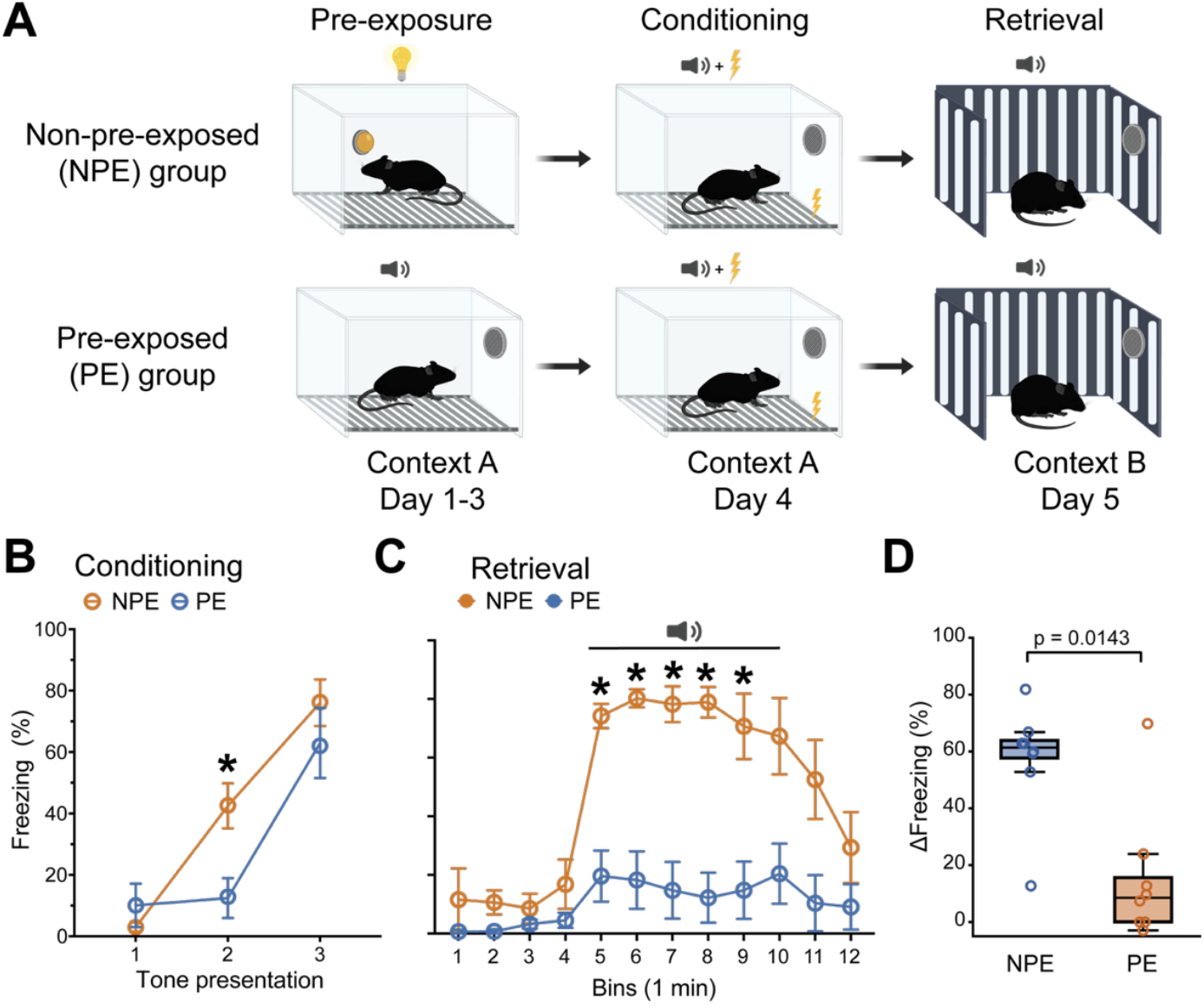
Pre-exposure to a Tone Attenuates Retrieval of a Tone-Evoked Conditioned Fear Response and Produces Latent Inhibition. **(A)** Schematic of the experimental design. Mice underwent a three-phase behavioral paradigm consisting of pre-exposure, conditioning, and retrieval. Pre-exposure and conditioning were conducted in Context A, which contained a grid floor, Plexiglass walls, and an anise odor. Retrieval was conducted in a distinct Context B, which contained a white floor, checkered or striped walls, and a vanilla odor. **(B)** Freezing during conditioning across tone–shock pairings. Freezing increased across tone presentations in both groups. Two-way repeated-measures ANOVA with Geisser– Greenhouse correction revealed a significant Tone × Pre-exposure interaction, F(1.92, 26.93) = 3.80, p = 0.0368 (generalized η^2^ = 0.128; medium-to-large effect size). Post hoc comparisons showed a significant difference between groups at the second tone presentation (*Šídák-corrected p = 0.0249), but not at the other tone presentations. **(C)** Freezing during retrieval, shown in 1-min time bins. NPE animals exhibited higher freezing than PE animals during tone presentation. Two-way repeated-measures ANOVA with Geisser–Greenhouse correction revealed a significant Time bin × Pre-exposure interaction, F(2.59, 36.29) = 6.44, p = 0.0020 (generalized η^2^ = 0.215; large effect size). Post hoc comparisons showed significant differences between groups only during tone presentation bins 1-5 (*Šídák-corrected p < 0.05). **(D)** ΔFreezing during tone presentation, calculated as the change in freezing between the 1-min period before tone onset and the 1-min period after tone onset. NPE animals displayed significantly greater ΔFreezing than PE animals, confirming that prior tone exposure produced latent inhibition (Mann– Whitney U = 9, two-tailed exact p = 0.0143; rank-biserial r = 0.719, large effect size). Data in **B** and **C** are shown as mean ± SEM. Data in **D** are shown as individual animals with median and interquartile range; whiskers indicate min–max. n = 8 mice per group (4 females and 4 males). See Table S1 for extended statistical results.

We first assessed fear acquisition during conditioning by measuring freezing during the 30-s tone presentations across tone–shock pairings (**Fig. 1B**). Freezing increased robustly across tone presentations, indicating successful acquisition of conditioned fear. Although the PE group showed lower freezing during the second tone presentation, both groups reached similar freezing levels by the third tone presentation. Thus, by the end of conditioning, PE and NPE mice showed indistinguishable conditioned responses, indicating that pre-exposure slowed the rate of fear acquisition but did not prevent conditioned fear learning.

We next examined conditioned responding during the retrieval session by analyzing freezing across 1-min time bins (**Fig. 1C**). NPE and PE animals responded differently during tone presentation: NPE mice showed sustained freezing throughout the tone, whereas PE mice showed very little freezing. These results indicate that NPE mice exhibited strong conditioned fear responses to the tone, while prior tone exposure selectively reduced conditioned responding during retrieval.

Because baseline freezing can vary across animals, and the initial moments of tone presentation typically evoke the strongest conditioned responses, we quantified tone-evoked freezing using Δfreezing, defined as the change in freezing from the 30 s before tone onset to the 30 s after tone onset. NPE animals showed significantly greater Δfreezing than PE animals, confirming that tone pre-exposure markedly attenuated conditioned responding (**Fig. 1D**). Together, these results demonstrate that prior exposure to the tone selectively reduces fear responding during retrieval without impairing acquisition, establishing a robust latent inhibition effect.

### Validation of Viral Targeting and Functional Efficacy of Chemogenetic Inhibition

To test the causal role of the LEC in latent inhibition during retrieval, we used a chemogenetic approach based on inhibitory Designer Receptors Exclusively Activated by Designer Drugs (DREADDs). To selectively manipulate LEC output neurons, we restricted viral expression of the inhibitory DREADD to glutamatergic neurons using the CaMKII promoter (AAV9-CaMKII-hM4Di-mCherry). Control mice received injections of the same virus expressing mCherry alone (**Fig. 2A**, *top*). All animals received clozapine- N-oxide (CNO, 1 mg/kg), the ligand used to activate the DREADD receptors, allowing temporally precise inhibition of LEC glutamatergic neurons during behavior in DREADD- expressing mice, but not in control mice.

**Figure 2.**
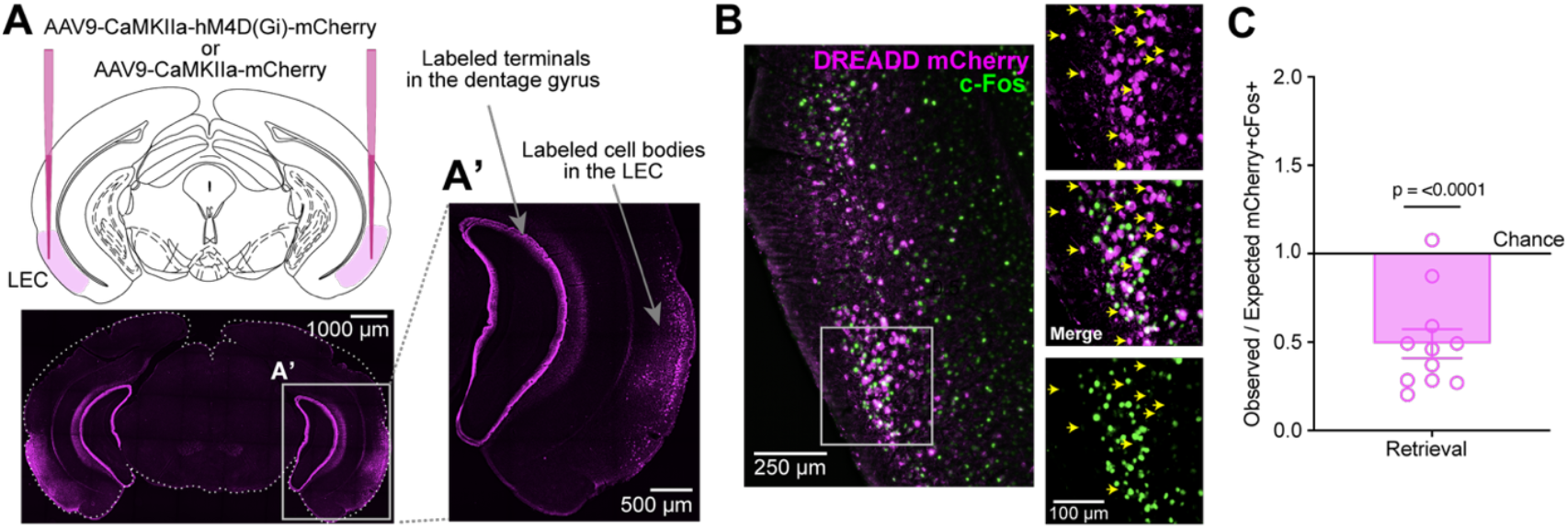
Validation of DREADD Expression in the LEC and Functional Efficacy of Chemogenetic Inhibition. **(A)** Schematic of viral injection targeting the lateral entorhinal cortex (LEC; top) and representative photomicrograph showing bilateral DREADD-mCherry expression in the LEC (bottom). **A’** Higher-magnification image showing mCherry immunoreactivity in LEC cell bodies and axonal projections to the dentate gyrus through the Perforant path, confirming selective labeling of LEC neurons. Animals lacking this expression pattern were excluded from the study. **(B)** Representative images showing DREADD-mCherry expression in magenta and c-Fos expression in green in the LEC. Insets on the right were taken from a region with high mCherry and c-Fos labeling and show mCherry^+^ cells (top), merged mCherry/c-Fos labeling (middle), and c-Fos^+^ cells (bottom). Yellow arrows indicate mCherry^+^ cells that do not express c-Fos, illustrating that even in a region with detectable c-Fos expression, most DREADD-expressing neurons were c-Fos negative and therefore not activated during retrieval following CNO treatment. **(C)** Quantification of observed mCherry^+^c-Fos^+^ colocalization relative to expected colocalization. Expected colocalization was calculated as:(*mcherry*^+^/*DAPI*^+^) × (*c*-*Fos*^+^/*DAPI*^+^) × *DAPI*^+^. An Observed/Expected MCherry+c-Fos+ colocalization ratio of 1 indicates colocalization at chance level. The observed/expected ratio was significantly below chance, indicating reduced activation of DREADD-expressing neurons following chemogenetic inhibition. One-sample t test versus theoretical mean = 1, *t*(10) = 6.247, *p* < 0.0001 (Cohen’s dz = −1.884, very large effect size). Data are presented as mean ± SEM.; n = 11 mice (4 females and 7 males).

To validate our viral targeting strategy, we verified both the restricted spread of viral expression within the LEC and its functional efficacy in producing chemogenetic inhibition. Only animals with bilateral LEC expression were included in subsequent analyses (**Fig. 2A**, *bottom*), and mice with bilateral spread into the hippocampus were excluded from the study. Bilateral expression of mCherry-tagged DREADDs was observed in LEC cell bodies and in axonal projections to the dentate gyrus, a major output region of the LEC, confirming accurate targeting of the injected region (**Fig. 2A’**).

To assess whether DREADD-expressing neurons were functionally inhibited, we quantified activity-dependent c-Fos expression within DREADD-mCherry-positive cells following CNO injection on the retrieval test day (**Fig. 2B**). Functional inhibition was assessed by comparing the observed colocalization of mCherry and c-Fos with the level expected by chance. Expected colocalization was calculated from the independent probabilities of cells being mCherry^+^ and c-Fos^+^ among DAPI^+^ nuclei, with DAPI^+^ nuclei used as the total cell count, and then multiplied by the total number of DAPI^+^ nuclei, as described previous ^18^. The observed/expected ratio of mCherry^+^/c-Fos^+^ colocalization was significantly below 1, the chance-predicted level (**Fig. 2C**), indicating that DREADD-expressing neurons were underrepresented among c-Fos^+^ activated cells following CNO administration. Together, these findings confirm both precise targeting of the LEC and the functional efficacy of chemogenetic inhibition, validating this approach for testing the role of the LEC in latent inhibition.

### Chemogenetic Inhibition of LEC During Retrieval Attenuates Latent Inhibition

We next asked whether LEC activity is required for the expression of latent inhibition. We hypothesized that, in tone-pre-exposed animals, the LEC contributes to retrieval of the prior experience in which the tone was inconsequential. This prior experience may interfere with conditioned fear responses when the same tone is later paired with shock, thereby reducing freezing during the retrieval test. To test this, we selectively inhibited LEC excitatory neurons during this session using inhibitory DREADDs. If the LEC supports access to the prior inconsequential-tone memory, then its inhibition should attenuate latent inhibition in PE mice, while leaving conditioned fear expression intact in NPE mice.

Mice were trained in the latent inhibition paradigm and received CNO only on Day 5 to selectively inhibit LEC neurons during the retrieval session (**Fig. 3A**). During conditioning, freezing increased across tone–shock pairings in both control (**Fig. 3B)** and DREADD-expressing (**Fig. 3E**) animals, with no differences between NPE and PE groups. In this cohort, we did not observe the reduced rate of acquisition seen in animals that did not receive viral injections (**Fig. 1B**). These results indicate that tone pre-exposure did not affect acquisition in this experiment and confirm that conditioned fear learning was intact across all groups before LEC inhibition.

**Figure 3.**
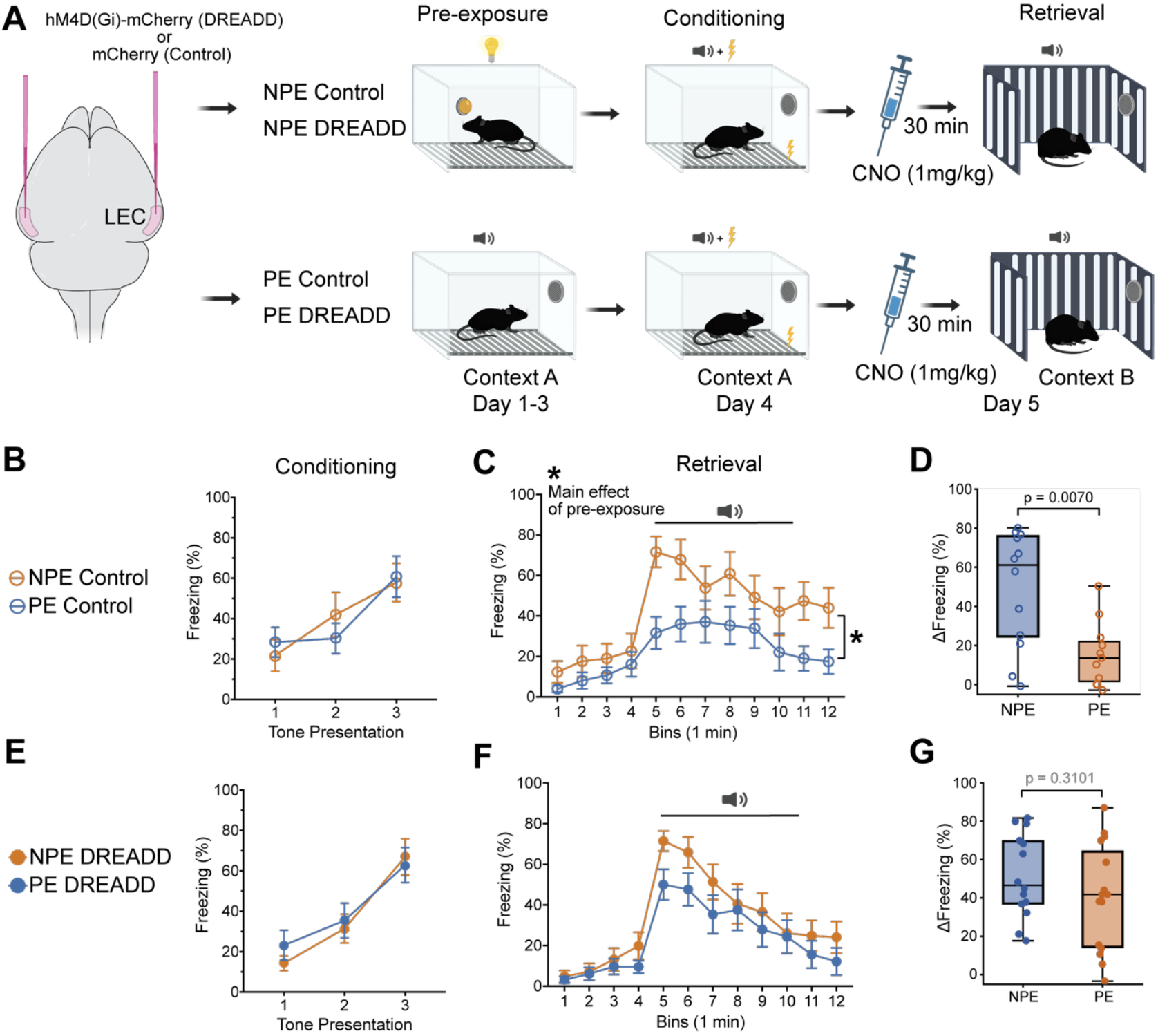
Chemogenetic inhibition of the LEC during retrieval attenuates latent inhibition. **(A)** Experimental design. Mice received bilateral LEC injections of AAV9-CaMKIIa-hM4Di-mCherry or AAV9-CaMKIIa-mCherry. Following pre-exposure and conditioning, CNO was administered 30 min before retrieval to inhibit LEC excitatory neurons in DREADD-expressing mice. **(B)** In Control animals, freezing increased across tone–shock pairings during conditioning, indicating intact acquisition. Two-way RM ANOVA with GG correction showed a significant main effect of Tone presentation, F(1.661, 33.21) = 10.91, p = 0.0005, generalized η^2^ = 0.192, with no main effect of Pre-exposure or Tone × Pre-exposure interaction. **(C)** During retrieval, Control animals showed higher freezing in NPE than PE mice, consistent with latent inhibition. Two-way RM ANOVA showed a significant main effect of Pre-exposure, F(1, 21) = 5.14, p = 0.0340, generalized η^2^ = 0.114, and a significant main effect of Time, with no Time × Pre-exposure interaction. **(D)** ΔFreezing was significantly reduced in PE compared with NPE Control mice, Mann–Whitney U = 23, p = 0.0070, rank-biserial r = 0.652. **(E)** In DREADD animals, freezing also increased across conditioning tones, F(1.86, 50.33) = 32.43, p < 0.0001, generalized η^2^ = 0.317, with no main effect of Pre-exposure or interaction. **(F)** During retrieval, DREADD animals showed no significant main effect of Pre-exposure, F(1, 27) = 1.68, p = 0.206, generalized η^2^ = 0.019, indicating attenuation of latent inhibition. **(G)** ΔFreezing did not differ between NPE and PE DREADD mice, Mann–Whitney U = 81, p = 0.310, rank-biserial r = 0.229. Data in **B, C, E**, and **F** are presented as mean ± SEM. Data in **D** and **G** show individual animals with median, interquartile range, and min–max whiskers. RM, repeated measures; GG, Geisser–Greenhouse. Group sizes were as follows: Control NPE, n = 12 mice, 5 females and 7 males; Control PE, n = 11 mice, 5 females and 6 males; DREADD NPE, n = 14 mice, 6 females and 8 males; and DREADD PE, n = 15 mice, 8 females and 7 males. See Table S1 for extended statistics.

We next examined the effect of inhibiting LEC CaMKIIα-expressing neurons during retrieval. To inhibit LEC neurons, mice received CNO (1 mg/kg) 30 min before the retrieval session. We then analyzed freezing across 1-min time bins in Control and DREADD-expressing animals (**Fig. 3C, F**). In control animals, freezing increased during tone presentation, with PE mice showing reduced freezing compared with NPE mice. In contrast, in DREADD-expressing animals, freezing increased during the tone in both the NPE and PE groups. To more sensitively capture tone-evoked responding at tone onset, we quantified Δfreezing. In control animals, PE mice exhibited significantly reduced Δfreezing compared with NPE mice, consistent with latent inhibition (**Fig. 3D**). In DREADD-expressing animals, however, there was no difference in ΔFreezing between PE and NPE mice (**Fig. 3G**), indicating that inhibition of the LEC during retrieval attenuated the expression of latent inhibition.

We also reanalyzed the data using a two-way ANOVA to directly compare Δfreezing between Control and DREADD groups as a function of pre-exposure. This analysis showed no significant Group × Pre-exposure interaction, F(1, 48) = 2.58, p = 0.115. Because our hypotheses specifically tested whether LEC inhibition affected tone- conditioned fear retrieval and/or latent inhibition, we performed 4 planned comparisons using the residual error term from the two-way ANOVA, with Šídák correction for 4 comparisons. Freezing did not differ between NPE Control and NPE DREADD animals, indicating that LEC inhibition did not impair retrieval of tone-conditioned fear responses (mean difference = −2.65%, t(48) = 0.27, p = 0.998). In Control animals, PE mice froze significantly less than NPE mice, confirming robust latent inhibition (mean difference = 33.41%, t(48) = 3.22, p = 0.009). In contrast, PE and NPE animals did not differ significantly in the DREADD group (mean difference = 11.12%, t(48) = 1.20, p = 0.656), suggesting that LEC inhibition attenuated the expression of latent inhibition. PE DREADD animals showed higher freezing than PE Control animals, but this comparison did not reach significance after correction for 4 planned comparisons (mean difference = −24.94%, t(48) = 2.53, p = 0.058).

Together, these findings indicate that LEC inhibition did not impair retrieval of tone- conditioned fear responses but attenuated the expression of latent inhibition. These results support a model in which the LEC contributes to retrieval-based memory interference.

## DISCUSSION

The present study showed that inhibition of LEC activity during retrieval attenuated latent inhibition without affecting the expression of conditioned fear. These findings suggest that the LEC supports the expression of prior stimulus experience during memory retrieval. More broadly, they are consistent with a role for the LEC in the competition between previously acquired and newly formed memory representations, providing evidence that the LEC contributes to memory interference.

Pre-exposed animals showed reduced freezing compared with non-pre-exposed controls during the retrieval test, as reflected in both time-bin analyses and Δfreezing measures. In our protocol, however, pre-exposure did not produce a consistent effect on the acquisition of fear conditioning. Although PE mice in the first experiment showed a slower rate of acquisition, this effect was not replicated in the second experiment. Moreover, in both experiments, freezing in PE mice did not differ from NPE mice by the third tone–shock pairing, indicating that both groups acquired the tone–shock association. Effects of stimulus pre-exposure on the acquisition of fear conditioning have been previously reported ^19–22^, and have been interpreted as evidence that latent inhibition reflects, at least in part, an acquisition deficit. This view forms the basis of acquisition/attentional theories, which propose that pre-exposure reduces attention to the stimulus and thereby slows its subsequent association with the unconditioned stimulus ^23–26^. In the present study, however, the largest and most consistent difference between NPE and PE animals emerged during retrieval rather than acquisition. Although reduced freezing in PE mice could reflect weaker conditioning, inhibition of the LEC revealed a latent fear response in pre-exposed animals, indicating that pre-exposure did not prevent acquisition of the tone–shock association. These findings support an interpretation in which latent inhibition, under the present experimental conditions, depends in part on retrieval-based interference. Specifically, retrieval of the prior memory that the tone was inconsequential may compete with retrieval or expression of the later tone–shock association, thereby reducing conditioned freezing. This interpretation is consistent with retrieval/competition models of latent inhibition ^6,8^.

Interestingly, recent evidence in Drosophila also supports a competition-based mechanism, in which reduced conditioned responding reflects competition between parallel memories formed during pre-exposure and conditioning ^27^.

The attenuation of latent inhibition following LEC inhibition suggests that this region supports memory interference during retrieval. Recent evidence indicates that activity in LEC ensembles can maintain episodic-memory representations necessary for recall ^28^. In the context of latent inhibition, the LEC may therefore preserve a representation of the pre-exposed tone as familiar and behaviorally irrelevant. However, this raises an important question: why is this representation not updated once the same tone is later paired with shock? One possibility is that the LEC does not receive sufficient modulatory input during conditioning to update its response to the tone. Ventral tegmental area dopamine neurons provide dense dopaminergic input to the LEC ^29^ and can update LEC responses to novel cues that predict an unconditioned stimulus ^12^. Dopamine neuron responses to novelty decrease with repeated stimulus exposure, and this repetition-dependent reduction is necessary for the expression of latent inhibition ^21,30^. Thus, reduced novelty-induced dopamine signaling in the LEC after pre-exposure may limit updating of the tone representation during conditioning, promoting interference during retrieval.

Furthermore, our results suggest that LEC activity may contribute to latent inhibition by suppressing the expression of the conditioned response, since inhibition of the LEC revealed a fear response in pre-exposed animals. The neural mechanism by which the LEC suppresses the expression of learned fear remains unknown. However, learned fear conditioning depends critically on the basolateral amygdala, whereas the expression of freezing depends on output pathways involving the central amygdala ^31^. Therefore, the LEC could influence latent inhibition by inhibiting central amygdala output, thereby reducing freezing during retrieval without preventing acquisition of the tone–shock association, which depends on the basolateral amygdala. Although the LEC projects to the central amygdala ^32^, the function of this pathway remains to be determined.

Some limitations should be considered. Chemogenetic inhibition was restricted to CaMKII-positive neurons, thereby targeting primarily excitatory neurons; however, DREADD expression was not layer-specific within the LEC. Because projections from deep LEC layers target the amygdala and may modulate fear responding, future studies using layer-specific targeting may reveal stronger or more selective effects. Finally, although the present findings support a role for the LEC during retrieval, we did not directly examine its contribution during the pre-exposure or conditioning phases.

In summary, the present findings support a role for the LEC in the expression of latent inhibition. More broadly, these results are consistent with the idea that the LEC contributes to retrieval-based memory interference by allowing prior experience to influence the expression of newly acquired associations. Given that the LEC is among the first regions affected during aging ^15,33^, LEC dysfunction may contribute to the worsening of memory interference with age.

## SUPPLEMENTAL INFORMATION

Table S1 with statistical results

## AUTHOR CONTRIBUTIONS

O.G contributed to conceptualization, investigation, methodology, data curation, formal analysis, visualization, and writing (original draft; review and editing). M.M

and J.N.T contributed to investigation and data curation. S.M. contributed to conceptualization, funding acquisition, supervision, and writing (review and editing).

## METHODS

### KEY RESOURCES TABLE

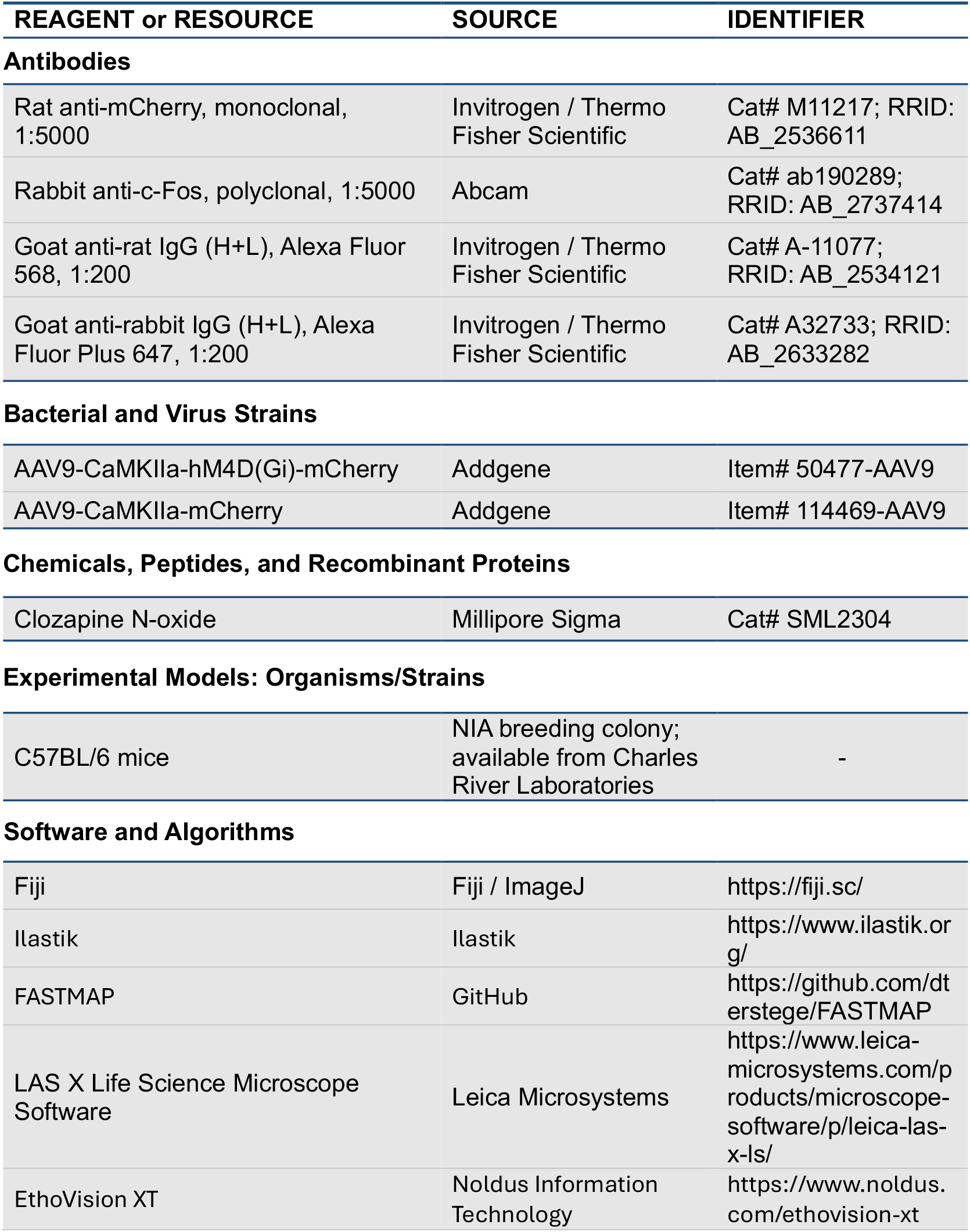

### LEAD CONTACT AND MATERIALS AVAILABILITY

Requests for additional information, resources, or reagents should be directed to the Lead Contact, Susana Mingote.

### EXPERIMENT MODEL AND SUBJECT DETAILS

#### Animals

Male and female wild-type C57BL/6 mice (4–7 months old) were co-housed in a temperature- and humidity-controlled facility on a reverse 12/12 h light/dark cycle (lights off at 11:00 AM), with ad libitum access to food, water, and enrichment materials. Mice were handled for 4 days (5 min) before the experiment to acclimate them to the experimenter. All experimental procedures were conducted during the dark phase, ensuring that behavioral testing was carried out under the dark cycle conditions. All protocols were approved by the Institutional Animal Care and Use Committee (IACUC) following NIH guidelines. All mice were obtained from C57BL/6 bred at the NIA and currently available from Charles River colonies.

## METHOD DETAILS

### Stereotaxic Injection of Viruses

Mice were anesthetized with isoflurane delivered by inhalation, with induction at 3% and maintenance at 1.5–3%, and placed in a Stoelting stereotaxic frame. Ophthalmic ointment containing neomycin and polymyxin B sulfates and bacitracin zinc was applied to prevent the eyes from drying during surgery. Small craniotomies were made above the target coordinates to allow viral injection into the LEC. Mice assigned to the DREADD group received bilateral injections of AAV9-CaMKIIa-hM4Di-mCherry, whereas control mice received AAV9-CaMKIIa-mCherry. Viral injections were performed using a Nanoject III injector (Drummond Scientific). The stereotaxic coordinates for LEC injections were AP −3.5 mm, ML ±4.4 to 4.5 mm, and DV −3.2 to −3.1 mm, with the injector positioned at a zero-degree angle. Mice were allowed at least two weeks for recovery and viral expression before behavioral testing.

### Immunohistochemistry and Imaging

Mice were deeply anaesthetized with ketamine and xylazine and were transcardially perfused with 1X phosphate-buffered saline (PBS) followed by 4%paraformaldehyde. Subsequently, brains were removed and stored overnight in 4% paraformaldehyde. Using a vibratome, the brains were sliced at 50-μm thickness. Slices were identified and mapped onto coronal sections of a mouse brain atlas (Paxinos and Franklin’s) and processed for immunohistochemistry staining. Tissue sections were washed three times in 1X PBS for 10 minutes each at room temperature to remove residual fixatives. To quench free aldehyde groups from prior fixation, slices were incubated in 0.1M glycine for 30 minutes, followed by three additional PBS washes. To minimize non-specific antibody binding, tissue was incubated in a blocking solution containing 10% normal goat serum (NGS) in 0.1% PBS-T (PBS with 0.1% Triton X-100) for 2 hours at room temperature.

For mCherry labeling, free-floating sections were incubated overnight at room temperature with rat anti-mCherry primary antibody (Invitrogen; 1:5000) diluted in blocking solution. The following day, sections were washed three times in PBS and incubated for 45 min at room temperature, protected from light, with Alexa Fluor 568- conjugated goat anti-rat secondary antibody diluted in 0.02% PBS-T. Sections were then washed three additional times in PBS, mounted onto glass slides in 0.1 M phosphate buffer, and coverslipped with ProLong Gold antifade reagent. Slides were allowed to dry overnight in a light-protected drawer before fluorescence imaging.

For animals processed for c-Fos immunohistochemistry, free-floating sections were incubated overnight with rat anti-mCherry primary antibody, as described above, together with rabbit anti-c-Fos primary antibody diluted in blocking solution. The following day, sections were washed in PBS and incubated for 45 min at room temperature, protected from light, with Alexa Fluor Plus 647-conjugated goat anti-rabbit secondary antibody diluted in 0.02% PBS-T. Sections were then washed, mounted, coverslipped, and imaged as described above.

mCherry and c-Fos fluorescence was visualized using a fluorescence microscope (Leica Microsystems, DM6B) to confirm viral targeting of the LEC. Only mice with confirmed viral expression in the correct anatomical region were included in the final analysis. For the DREADD-injected group, mice were excluded if mCherry expression was restricted to only one hemisphere, absent from the bilateral LEC, or observed bilaterally in the hippocampus rather than the intended LEC target.

### Drug Preparation and Injection

Clozapine-N-Oxide (CNO) was diluted in saline (0.9% sodium chloride) and injected intraperitoneal at a volume of 10 ml/kg for a final concentration of 1mg/kg.

To minimize potential stress-related confounds associated with intraperitoneal injections, a subset of animals was habituated to the injection procedure by receiving daily intraperitoneal (i.p.) injections of sterile saline (0.9%) throughout the behavioral protocol. On the retrieval test day, saline was replaced with clozapine-N-oxide (CNO; 1 mg/kg, i.p.), allowing chemogenetic inhibition to be performed in injection-habituated animals. Animals that were not habituated to injections received CNO only on the retrieval day. Data from injection-habituated and non-habituated animals were pooled for analysis because no observable behavioral differences were detected between these groups within the DREADD cohort.

### Behavioral Apparatus

Behavioral testing was conducted in four identical conditioning chambers (Noldus) including sound attenuating cases. The inner chambers consisted of a plain Plexiglas box with a Plexiglas door. The floor of the inner conditioning chamber is comprised of a shock delivery system, consisting of steel bars. The chambers were equipped with overhead white lights and a speaker for tone presentations. Freezing behavior was recorded using an automated tracking system (EthoVision).

### Latent Inhibition Paradigm

To establish latent inhibition, mice were assigned to either a pre-exposed (PE) or non-pre-exposed (NPE) group. **Pre-exposure phase (Day 1-3):** PE mice received 20 presentations per session of a1400 Hz, 80 dB tone (30 s duration) across three consecutive days. Stimuli were delivered at variable interstimulus intervals (30–90 s) to prevent temporal predictability. NPE mice received an equivalent number of light stimuli and were not exposed to the tone. Pre-exposure was conducted in the fear conditioning chambers. Pre-exposure was conducted in the fear conditioning box described above. An anise odor was added to the context (context A). **Conditioning (Day 4):** All mice underwent fear conditioning in the same context as pre-exposure (Context A). The tone (30 s) co-terminated with a 2 s foot shock (0.7 mA), and this pairing was repeated three times. The intensity of the shock was verified using an Ammeter (Med Associates Inc, ENV-421). **Recall test (Day 5):** Testing was conducted in a distinct context (Context B), characterized by a vanilla scent, checkered-pattern walls, a striped back panel, and a white plastic floor, to minimize contextual fear generalization. Mice were first exposed to a 4-min baseline period, followed by a 6-min continuous tone presentation to assess cue-evoked freezing behavior. All animals in both control and DREADD groups received an intraperitoneal injection of Clozapine-N-oxide (CNO; 1 mg/kg) 30 min prior to behavioral testing. A subset of animals was subsequently perfused 1–1.5 h following the onset of the behavioral session to assess c-Fos expression associated with recall-related neural activity.

## QUANTIFICATION AND STATISTICAL ANALYSIS

### Quantification and Analysis of Immunohistochemistry

For quantification of labeling, c-Fos^+^ cells were identified using Ilastik pixel classification to generate binary images (Berg et al., 2019), and cell counts were extracted from regions of interest using FASTMAP (Terstege et al., 2022). mCherry^+^ DREADD-expressing cells were quantified manually in ImageJ using the multipoint tool. Cells were counted in two brain sections per animal, corresponding to approximately −3.5 and −3.6 mm from bregma. For each animal, colocalization values were averaged across the two sections. For histological validation, the probability of colocalization was calculated as the quantification of observed mCherry^+^c-Fos^+^ colocalization relative to the expected colocalization. The expected colocalization was calculated as (mCherry^+^/DAPI^+^) × (c-Fos^+^/DAPI^+^) × DAPI^+^ . The ratio of observed and expected colocalization was compared to a theoretical mean of 1 using a one-sample t-test.

### Statistical Analysis of Behavioral Data

Behavioral data were analyzed using GraphPad Prism version 11.0.0. Freezing was expressed as the percentage of time spent immobile. Sex differences in Δfreezing were assessed within each treatment/pre-exposure group using Welch’s t-tests. Because no significant differences were detected between males and females, data were combined across sex.

For all repeated-measures ANOVAs, the Geisser–Greenhouse correction was applied to within-subject effects and interactions involving within-subject factors, and corrected degrees of freedom are reported. During conditioning and retrieval, freezing was analyzed using repeated-measures two-way ANOVA, with time, defined as tone presentations or time bins, as the within-subject factor and pre-exposure as the between-subject factor. When a significant interaction was detected, effects were followed by Šídák-corrected post hoc comparisons, when applicable. For analyses in which the interaction was not significant but specific a priori hypotheses were tested, planned comparisons were performed using the residual error term from the two-way ANOVA model. P values from these planned comparisons were adjusted using the Šídák correction for the four predefined comparisons: NPE Control versus NPE DREADD, PE Control versus PE DREADD, NPE Control versus PE Control, and NPE DREADD versus PE DREADD.

To more sensitively quantify tone-evoked responding, Δfreezing was calculated as: freezing during the 1 min after tone onset minus freezing during the 1 min immediately before tone onset. Because Δfreezing data were not consistently normally distributed across experiments, comparisons between two independent groups were analyzed using Mann–Whitney U tests. Δfreezing data are shown as individual animals overlaid on box-and-whisker plots, with boxes indicating the interquartile range, center lines indicating the median, and whiskers indicating the minimum and maximum values.

Assumptions of normality were assessed using Shapiro–Wilk tests, and equality of variance was assessed using F tests. Statistical significance was set at p < 0.05. Effect sizes were reported where applicable. Data are presented as mean ± SEM.

For a detailed description of the statistical analyses used for each figure, see Table S1.

---

**Supplemental Table S1.**
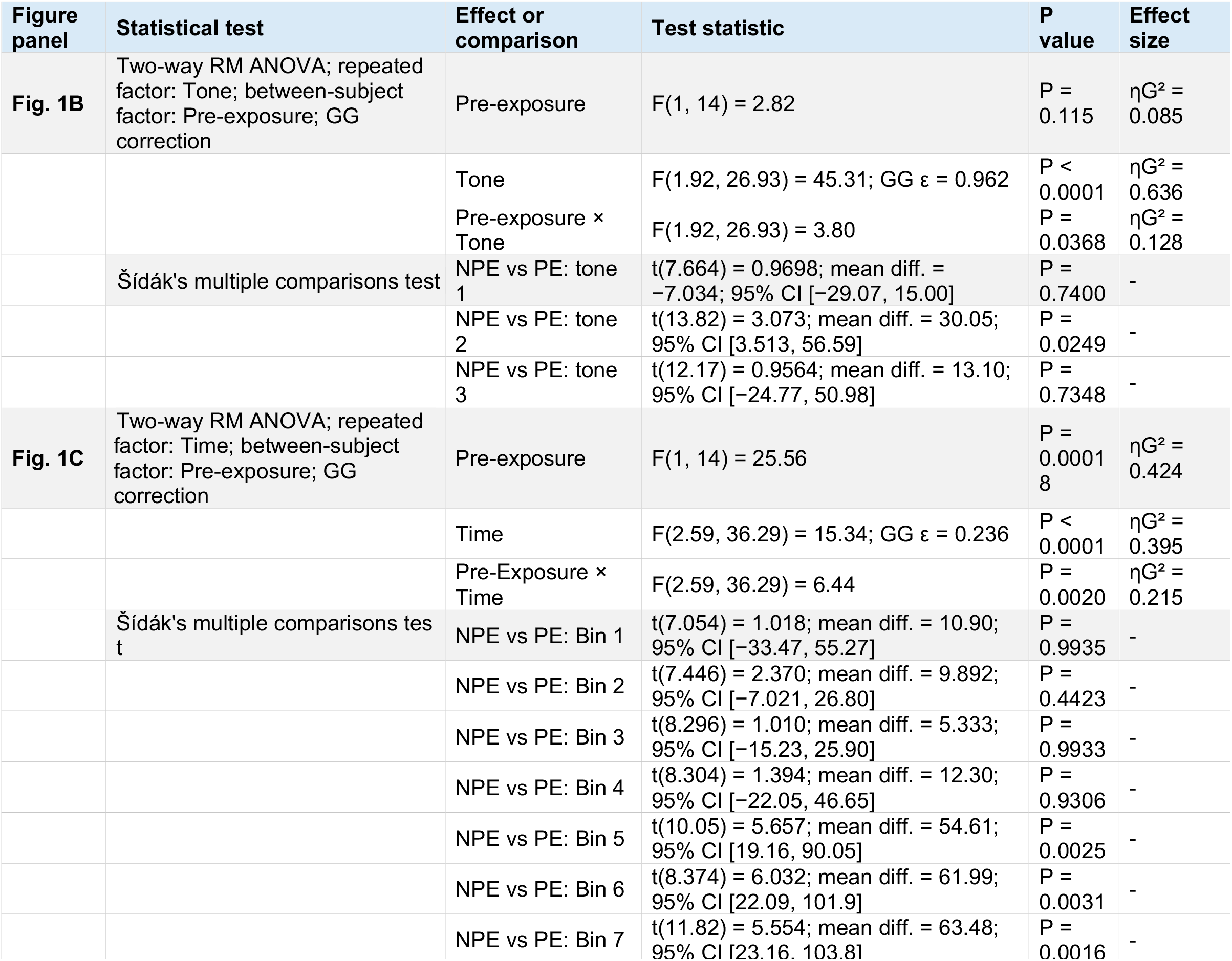
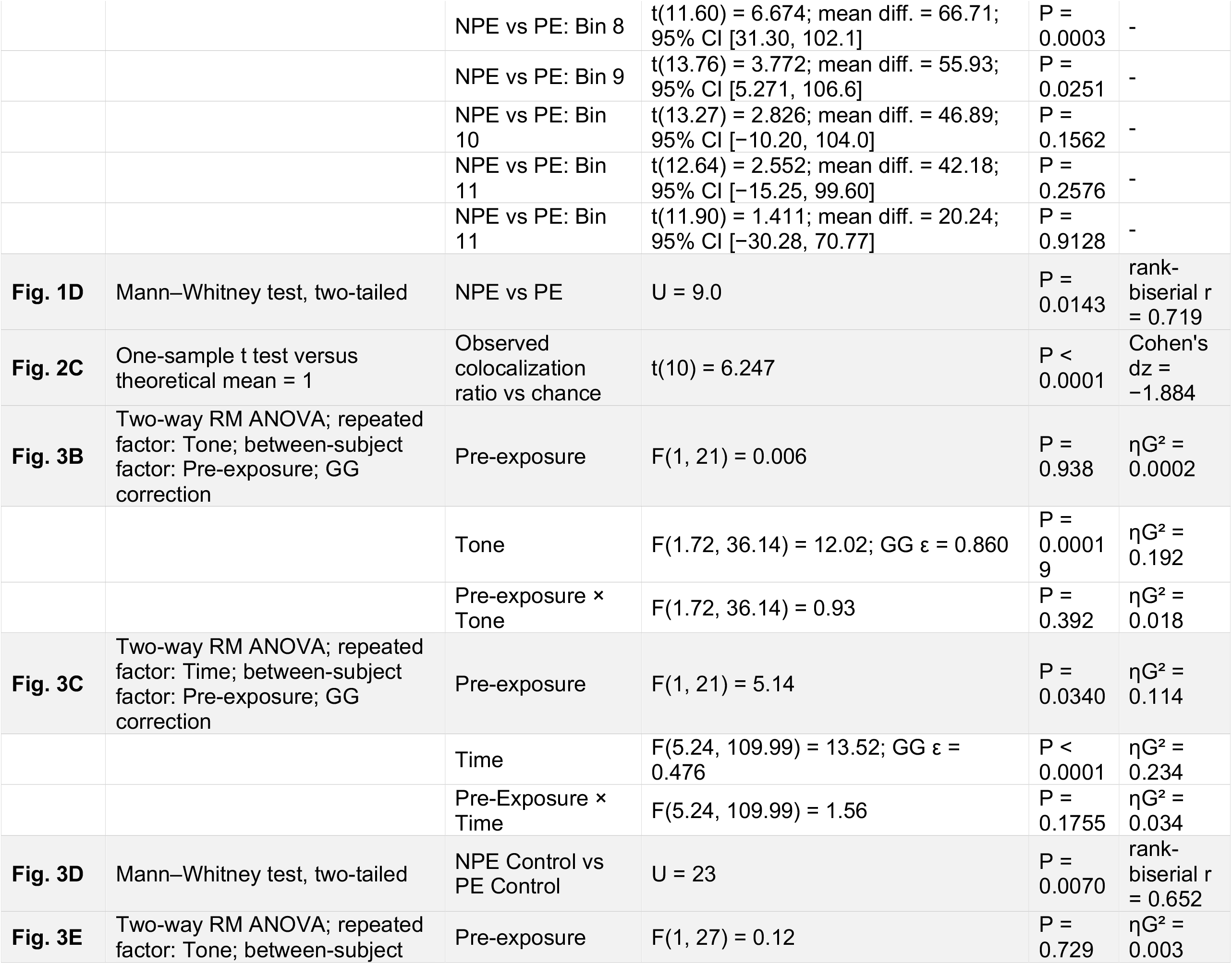
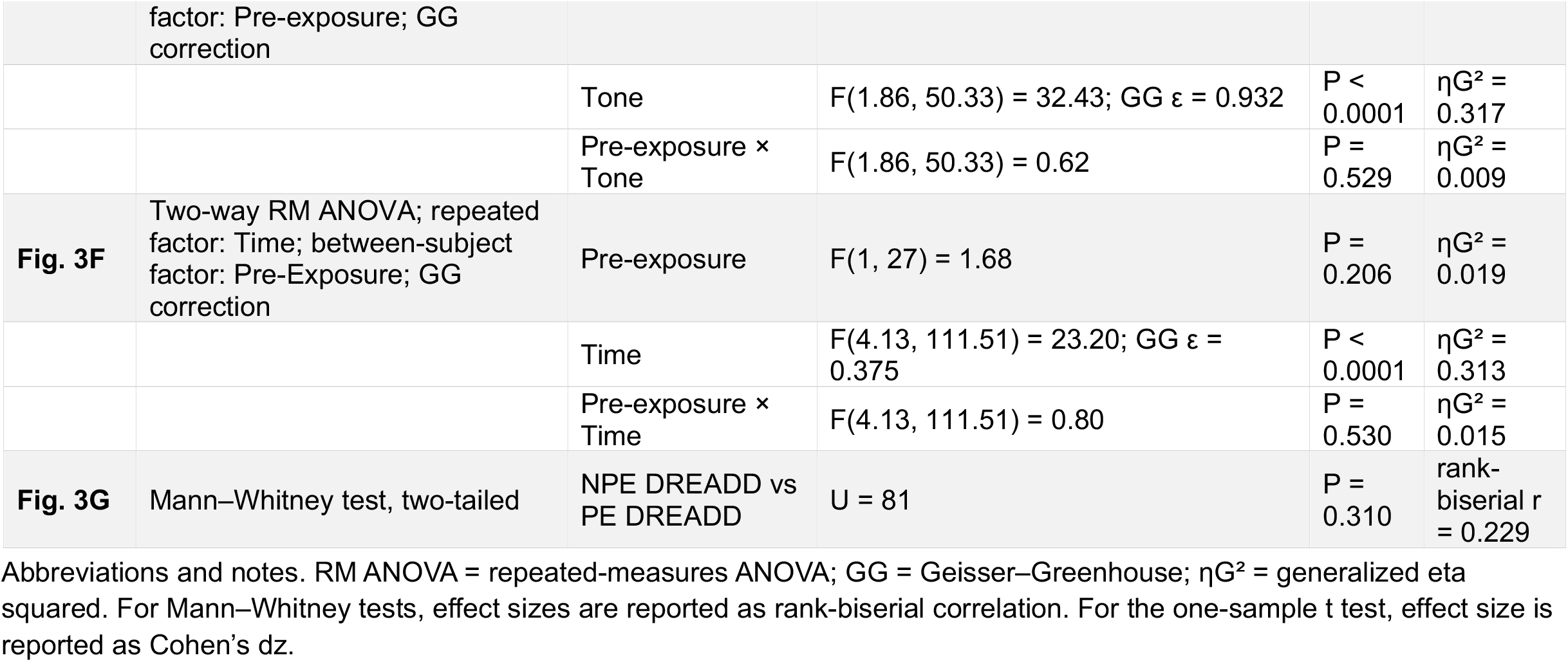
Statistical analyses and effect sizes for Figures 1–3.

