## Supplemental Table S1 for "Dissecting the Role of the Lateral Entorhinal Cortex in Memory Interference"

**Supplemental Table S1. Statistical analyses and effect sizes for Figures 1–3**

| Figure panel | Statistical test | Effect or comparison | Test statistic | P value | Effect size |
|---|---|---|---|---|---|
| **Fig. 1B** | Two-way RM ANOVA; repeated factor: Tone; between-subject factor: Pre-exposure; GG correction | Pre-exposure | $F(1, 14) = 2.82$ | $P = 0.115$ | $\eta G^2 = 0.085$ |
| | | Tone | $F(1.92, 26.93) = 45.31$; GG $\varepsilon = 0.962$ | $P < 0.0001$ | $\eta G^2 = 0.636$ |
| | | Pre-exposure × Tone | $F(1.92, 26.93) = 3.80$ | $P = 0.0368$ | $\eta G^2 = 0.128$ |
| | Šídák's multiple comparisons test | NPE vs PE: tone 1 | $t(7.664) = 0.9698$; mean diff. = −7.034; 95% CI [−29.07, 15.00] | $P = 0.7400$ | - |
| | | NPE vs PE: tone 2 | $t(13.82) = 3.073$; mean diff. = 30.05; 95% CI [3.513, 56.59] | $P = 0.0249$ | - |
| | | NPE vs PE: tone 3 | $t(12.17) = 0.9564$; mean diff. = 13.10; 95% CI [−24.77, 50.98] | $P = 0.7348$ | - |
| **Fig. 1C** | Two-way RM ANOVA; repeated factor: Time; between-subject factor: Pre-exposure; GG correction | Pre-exposure | $F(1, 14) = 25.56$ | $P = 0.0001$ 8 | $\eta G^2 = 0.424$ |
| | | Time | $F(2.59, 36.29) = 15.34$; GG $\varepsilon = 0.236$ | $P < 0.0001$ | $\eta G^2 = 0.395$ |
| | | Pre-Exposure × Time | $F(2.59, 36.29) = 6.44$ | $P = 0.0020$ | $\eta G^2 = 0.215$ |
| | Šídák's multiple comparisons tes t | NPE vs PE: Bin 1 | $t(7.054) = 1.018$; mean diff. = 10.90; 95% CI [−33.47, 55.27] | $P = 0.9935$ | - |
| | | NPE vs PE: Bin 2 | $t(7.446) = 2.370$; mean diff. = 9.892; 95% CI [−7.021, 26.80] | $P = 0.4423$ | - |
| | | NPE vs PE: Bin 3 | $t(8.296) = 1.010$; mean diff. = 5.333; 95% CI [−15.23, 25.90] | $P = 0.9933$ | - |
| | | NPE vs PE: Bin 4 | $t(8.304) = 1.394$; mean diff. = 12.30; 95% CI [−22.05, 46.65] | $P = 0.9306$ | - |
| | | NPE vs PE: Bin 5 | $t(10.05) = 5.657$; mean diff. = 54.61; 95% CI [19.16, 90.05] | $P = 0.0025$ | - |
| | | NPE vs PE: Bin 6 | $t(8.374) = 6.032$; mean diff. = 61.99; 95% CI [22.09, 101.9] | $P = 0.0031$ | - |
| | | NPE vs PE: Bin 7 | $t(11.82) = 5.554$; mean diff. = 63.48; 95% CI [23.16, 103.8] | $P = 0.0016$ | - |

| Figure panel | Statistical test | Effect or comparison | Test statistic | P value | Effect size |
|---|---|---|---|---|---|
| | | NPE vs PE: Bin 8 | t(11.60) = 6.674; mean diff. = 66.71; 95% CI [31.30, 102.1] | P = 0.0003 | - |
| | | NPE vs PE: Bin 9 | t(13.76) = 3.772; mean diff. = 55.93; 95% CI [5.271, 106.6] | P = 0.0251 | - |
| | | NPE vs PE: Bin 10 | t(13.27) = 2.826; mean diff. = 46.89; 95% CI [−10.20, 104.0] | P = 0.1562 | - |
| | | NPE vs PE: Bin 11 | t(12.64) = 2.552; mean diff. = 42.18; 95% CI [−15.25, 99.60] | P = 0.2576 | - |
| | | NPE vs PE: Bin 11 | t(11.90) = 1.411; mean diff. = 20.24; 95% CI [−30.28, 70.77] | P = 0.9128 | - |
| **Fig. 1D** | Mann–Whitney test, two-tailed | NPE vs PE | U = 9.0 | P = 0.0143 | rank-biserial r = 0.719 |
| **Fig. 2C** | One-sample t test versus theoretical mean = 1 | Observed colocalization ratio vs chance | t(10) = 6.247 | P < 0.0001 | Cohen's dz = −1.884 |
| **Fig. 3B** | Two-way RM ANOVA; repeated factor: Tone; between-subject factor: Pre-exposure; GG correction | Pre-exposure | F(1, 21) = 0.006 | P = 0.938 | ηG² = 0.0002 |
| | | Tone | F(1.72, 36.14) = 12.02; GG ε = 0.860 | P = 0.00019 | ηG² = 0.192 |
| | | Pre-exposure × Tone | F(1.72, 36.14) = 0.93 | P = 0.392 | ηG² = 0.018 |
| **Fig. 3C** | Two-way RM ANOVA; repeated factor: Time; between-subject factor: Pre-exposure; GG correction | Pre-exposure | F(1, 21) = 5.14 | P = 0.0340 | ηG² = 0.114 |
| | | Time | F(5.24, 109.99) = 13.52; GG ε = 0.476 | P < 0.0001 | ηG² = 0.234 |
| | | Pre-Exposure × Time | F(5.24, 109.99) = 1.56 | P = 0.1755 | ηG² = 0.034 |
| **Fig. 3D** | Mann–Whitney test, two-tailed | NPE Control vs PE Control | U = 23 | P = 0.0070 | rank-biserial r = 0.652 |
| **Fig. 3E** | Two-way RM ANOVA; repeated factor: Tone; between-subject | Pre-exposure | F(1, 27) = 0.12 | P = 0.729 | ηG² = 0.003 |

| Figure panel | Statistical test | Effect or comparison | Test statistic | P value | Effect size |
|---|---|---|---|---|---|
| | factor: Pre-exposure; GG correction | | | | |
| | | Tone | F(1.86, 50.33) = 32.43; GG ε = 0.932 | P < 0.0001 | ηG² = 0.317 |
| | | Pre-exposure × Tone | F(1.86, 50.33) = 0.62 | P = 0.529 | ηG² = 0.009 |
| **Fig. 3F** | Two-way RM ANOVA; repeated factor: Time; between-subject factor: Pre-Exposure; GG correction | Pre-exposure | F(1, 27) = 1.68 | P = 0.206 | ηG² = 0.019 |
| | | Time | F(4.13, 111.51) = 23.20; GG ε = 0.375 | P < 0.0001 | ηG² = 0.313 |
| | | Pre-exposure × Time | F(4.13, 111.51) = 0.80 | P = 0.530 | ηG² = 0.015 |
| **Fig. 3G** | Mann–Whitney test, two-tailed | NPE DREADD vs PE DREADD | U = 81 | P = 0.310 | rank-biserial r = 0.229 |

Abbreviations and notes. RM ANOVA = repeated-measures ANOVA; GG = Geisser–Greenhouse; ηG² = generalized eta squared. For Mann–Whitney tests, effect sizes are reported as rank-biserial correlation. For the one-sample t test, effect size is reported as Cohen’s dz.
